# MODULATION OF ARCHAEAL HYPERNUCLEOSOME STRUCTURE AND STABILITY BY Mg^2+^

**DOI:** 10.1101/2025.05.26.656202

**Authors:** Ilias Zarguit, Marc K. M. Cajili, Bert van Erp, Samuel Schwab, Nico van der Vis, Marianne Bakker, John van Noort, Remus T. Dame

**Affiliations:** Leiden Institute of Chemistry, Leiden University, Einsteinweg 55, 2333CC Leiden, The Netherlands; Centre for Microbial Cell Biology, Leiden University, Leiden, The Netherlands; Centre for Interdisciplinary Genome Research, Leiden University, Leiden, The Netherlands; Leiden Institute of Physics, Leiden University, Einsteinweg 55, 2333CC, Leiden, The Netherlands

## Abstract

DNA-wrapping histone proteins play a central role in chromatin organization, gene expression and regulation in most eukaryotes and archaea. While the structure and function of eukaryotic histones are well-characterized, archaeal histones and their complexes with DNA require further scrutiny. Distinct from their eukaryotic counterparts, previously characterized canonical archaeal histones assemble on DNA into an ‘endless’ superhelical nucleoprotein complex called a hypernucleosome. In this study, we explored whether hypernucleosome formation is a conserved feature of canonical archaeal histones. Moreover, to further elucidate how hypernucleosomes are regulated, we also explored how changes in the physico-chemical conditions, particularly the presence of Mg^2+^, influence the hypernucleosome. Using a combination of Tethered Particle Motion (TPM) and single-molecule force spectroscopy, we established that *T. kodakarensis* histones assemble into hypernucleosomes on DNA, similar to the *M. fervidus* histones HMfA and HMfB, the only canonical histones structurally characterized in previous studies. However, the effects of Mg^2+^ ions are distinct despite the histones’ high sequence- and structural similarity. We propose a model in which Mg^2+^ ions exert a generic effect on hypernucleosome compactness and stability due to electrostatic DNA shielding, with additional differential effects depending on histone identity.

## INTRODUCTION

Organisms dynamically organize their genomes, structurally and functionally, in response to changes in environmental conditions, cell cycle stages, and DNA transactions. Typically, architectural proteins, like the nucleoid-associated proteins (NAPs) in bacteria (1-4), mitochondria (5), and some archaea (6-8) facilitate this process by bending or bridging DNA (9). All eukaryotes and most archaea encode histone proteins (6,10-14) that feature a signature motif of 3 alpha helices linked by two loops, known as the histone fold. Histones assemble into a multimeric protein core that wraps DNA to form nucleosomes (15). Recently, our group identified many different classes of atypical histones (14). Among these, some histones showed a gain of function due to of an N-terminal extension, which facilitates the formation of histone tetramers that bridge DNA (14,16), similar to many NAPs. Based on structure, histone proteins were also identified in some bacteria (17-20). While in one of the studies it was proposed that the bacterial histone HBb coats and straightens DNA (19), a subsequent study suggested that these proteins bind and bend DNA as dimers (18).

Archaeal histones form nucleosomes distinct from their eukaryotic counterparts. While eukaryotic histones assemble into an octameric histone core of two H2A-H2B histone dimers and one H3-H4 histone tetramer that is wrapped by around 147 base pairs of DNA, yielding the canonical eukaryotic nucleosome (15), archaeal histones were originally suggested to form a tetrameric histone core composed of histone homodimers or heterodimers (21). MNase experiments supported this tetrameric nucleosome model in *Methanothermus fervidus* (22), showing protection of 60 bp DNA fragments. SELEX-optimized high-affinity sites yielded size-delimited structures and nucleosomes with a tetrameric histone core (23,24). However, studies on *Thermococcus kodakarensis* chromatin suggested a more complex picture: histones might form extended DNA-wrapping multimers wherein histone dimers, each binding to a minimal 30 base pairs DNA, cluster together (25,26). X-ray crystallography studies on HMfB-DNA complexes have revealed that, indeed, DNA is wrapped around archaeal histones in a continuous superhelix (27), forming a structure known as a hypernucleosome (28). Stacking these histone dimers brings adjacently positioned lysines and glutamic acids together, forming electrostatic interactions (referred to as ‘stacking interactions’) that potentially stabilize the hypernucleosome complex (10). Mechanical characterization of hypernucleosomes has established the crucial importance of stacking interaction residues in obtaining a compact and mechanically stable structure (28).

Modulation of chromatin structure and gene accessibility by histones results in a dynamic, environmentally responsive genome crucial for cell survival, adaptation, and evolution. While complex ATP-dependent remodelling machinery has been shown to modulate chromatin structure in eukaryotes (29), there is no evidence of similar mechanisms in archaea to date. Instead, proposed mechanisms for hypernucleosome modulation include binding of polymerization-deficient histone variants acting as capstones (11), specific sequences limiting hypernucleosome growth (24), and post-translational modifications at the stacking interface that may alter hypernucleosome structure and stability (30).

Similar to bacterial chromosomes, archaeal chromosomes may act as sensors of environmental changes, such as changes in K^+^ or Mg^2+^ concentration, altering local chromatin structure, thus affecting gene activity (31-33). Generic ionic effects are due to shielding of surface charge on proteins or DNA (34), affecting protein-DNA, protein-protein, or DNA-DNA interactions (35), affecting eukaryotic nucleosome assembly and nucleosome-nucleosome interactions (36). In eukaryotic histones, it was observed that the presence of Mg^2+^ can induce self-association of oligonucleosomes, leading to a more compact structure (37). In the presence of Mg^2+^, short 207 base pair hypernucleosomes of archaeal histone HTkC become more compact (38). Single-particle cryogenic electron microscopy studies revealed the concurrent existence of stacked and kinked conformations of such short hypernucleosomes in the presence of Mg^2+^ (38). However, it remains unclear how archaeal histones and hypernucleosomes respond to Mg^2+^, possibly regulating gene expression.

In this study, we employ tethered particle motion (TPM) analysis (39) and single-molecule force spectroscopy (40) to investigate the formation of hypernucleosomes on DNA by recombinant histones HTkA and HTkB from *Thermococcus kodakarensis*, as well as histones HMfA and HMfB from *Methanothermus fervidus*. We demonstrate that HTk histones in complex with DNA yield hypernucleosomes, confirming observations for HMf histones (28). Additionally, we show that Mg^2+^ influences hypernucleosome structure and stability, with distinct differences exhibited by the histone variants.

## RESULTS

### HTkA and HTkB histones from *T. kodakarensis* form hypernucleosomes on DNA

To investigate whether histones HTkA and HTkB assemble into hypernucleosomes, we conducted TPM experiments with a linear 685 base pair fragment. TPM is a single-molecule technique that allows for the observation of protein-DNA interactions. In brief, a polystyrene bead is tethered to a glass surface with a DNA duplex acting as a protein-binding substrate. The motion of the tethered DNA serves as a representation of DNA conformation. Any change in the conformation, measured by a change in the root mean squared (RMS) displacement of the bead, can then be described as a function of the protein concentration. An increase in HTkB concentration yields a reduction in RMS from about 160 nm to 75 nm, with protein saturation being reached at 10 nM (Figure 1A). An increase in HTkA concentration yields a reduction in RMS to 80 nm, indicating that both histones compact DNA to a similar extent (Figure 1A). The observed decrease in RMS is similar to that observed in experiments with HMfA and HMfB, indicating DNA wrapping and hypernucleosome formation (28). To quantitatively compare different histone proteins, we determined the ‘midpoint concentration’, which is defined as the protein concentration at which the RMS is exactly in between that of the fully unbound and saturated states. The midpoint concentration for HTkB (1.33 ± 1.1 nM) was lower than that for HTkA (9.21 ± 1.1 nM) (see Table 1), indicative of a higher DNA binding affinity for HTkB. Alanine substitution of the lysine residues predicted to be involved in electrostatic stacking in HTkB_K27A,K62A,K66A_ (10) yielded an increase in the midpoint concentration to 271.8 ± 1.1 nM (Supplementary Figure 2), higher than the wildtype, and in line with earlier experiments on similar variants of HMfA and HMfB (28). We also estimated the end-to-end distance (EED) of individual DNA tethers as a function of protein concentration. The EED at saturation is 33 ± 3 nm for HTkA and 26 ± 8 nm for HTkB (Supplementary Figure 3 and Table 1), consistent with the EED expected based on the HMfB-DNA co-crystal structure (27).

**Table 1.** An overview of RMS midpoint concentrations, RMS and end-to-end distances (at saturating protein concentration) in absence and presence of 5 mM Mg^2+^.

|  | HTkA |  | HTkB |  | HMfA |  | HMfB |  |
| --- | --- | --- | --- | --- | --- | --- | --- | --- |
| | 0 mM $Mg^{2+}$ | 5 mM $Mg^{2+}$ | 0 mM $Mg^{2+}$ | 5 mM $Mg^{2+}$ | 0 mM $Mg^{2+}$ | 5 mM $Mg^{2+}$ | 0 mM $Mg^{2+}$ | 5 mM $Mg^{2+}$ |
| Midpoint concentration (nM) | $9.2 \pm 1.1$ | $1.3 \pm 1.1$ | $2.5 \pm 1.1$ | $1.8 \pm 1.1$ | $59.7 \pm 1.2$ | $11.6 \pm 1.1$ | $9.0 \pm 1.0$ | $7.3 \pm 1.1$ |
| RMS (nm) | $80 \pm 4$ | $69 \pm 5$ | $75 \pm 4$ | $64 \pm 5$ | $86 \pm 5$ | $83.1 \pm 2$ | $80 \pm 3$ | $72 \pm 8$ |
| End-to-end distance (nm) | $33 \pm 3$ | $22 \pm 11$ | $26 \pm 8$ | $19 \pm 9$ | $33 \pm 5$ | $35 \pm 12$ | $22 \pm 15$ | $18 \pm 10$ |

**Figure 1.**
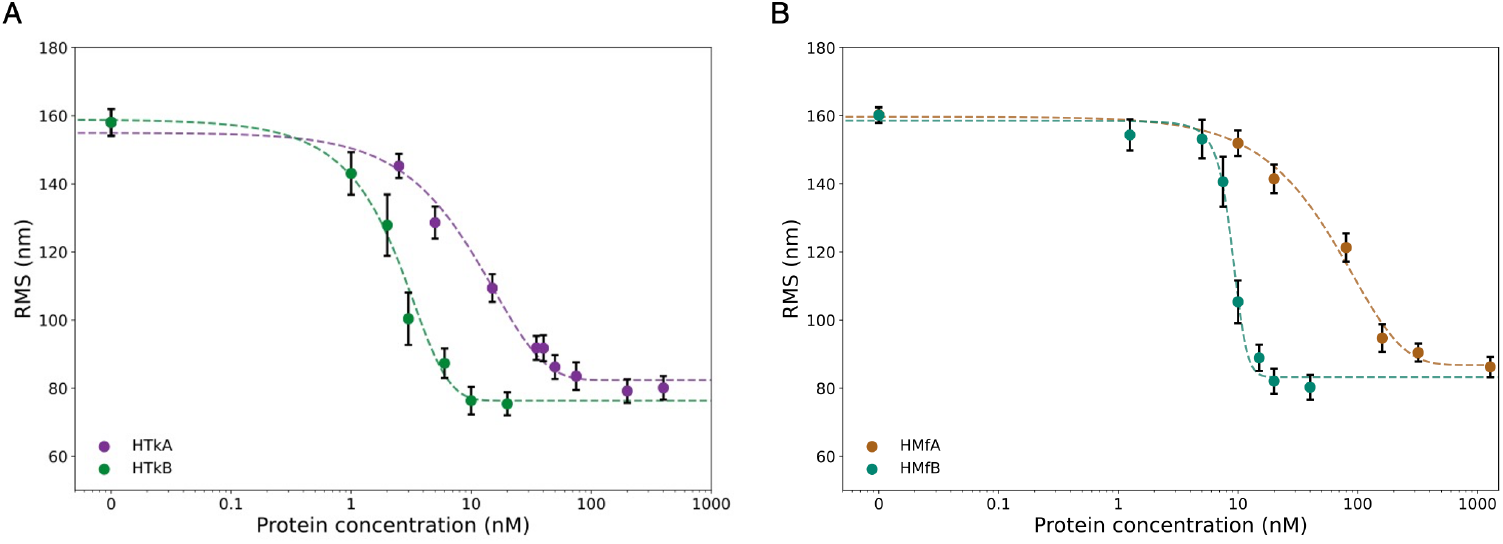
T. kodakarensis and M. fervidus histones form hypernucleosomes upon binding DNA. TPM experiments show that (A) HTkA and HTkB and (B) HMfA and HMfB assemble into hypemucleosomes on DNA substrates in 25 mM Tris-HCl pH 7 and 75 mM KC1. Each measurement (N > 100) was performed in duplicate, with the error bars representing the standard deviation. Dashed lines are added to guide the eye.

Next, we repeated earlier TPM experiments on histones HMfA and HMfB from *M. fervidus* (28) to directly compare their DNA binding properties with the histones from *T. kodakarensis*. For both proteins, a reduction in RMS was observed at saturating protein concentration in line with hypernucleosome formation (Figure 1B), confirming earlier data (28). Yet, the two proteins differed by almost an order of magnitude in midpoint concentration (9.0 ± 1.0 nM for HMfB vs 59.7 ± 1.2 nM for HMfA), with much lower protein concentration required to achieve compaction by HMfB (Table 1). Furthermore, the slope at the midpoint is larger for HMfA than for HMfB (Figure 1B). Together these observations indicate that HMfB has a higher DNA affinity than HMfA and is possibly more cooperative in binding DNA than HMfA, consistent with previous observations (24).

### Mg^2+^ ions promote hypernucleosome formation and DNA compaction by archaeal histones

Previous studies using sedimentation velocity analytical ultracentrifugation (SV-AUC) (38) demonstrated that Mg^2+^ influences compaction of short hypernucleosomes, accommodating 4 to 5 histone dimers. These experiments were limited to short DNA substrates and performed with HTkA histones only. Here, we study the effect of Mg^2+^ on substrates allowing binding of an order of magnitude more histone dimers. To study the universality of Mg^2+^ effects on hypernucleosome complexes from other species, we performed TPM experiments on HTkA/B and HMfA/B histones in the presence and absence of Mg^2+^. Results indicate that Mg^2+^ promotes hypernucleosome formation and enhances compaction for both HTk and HMf histones (Figure 2 and Table 1), in agreement with previous studies showing increased compaction in the presence of Mg^2+^ (38). Notably, that both HTkA and HMfA exhibited a stronger response to Mg^2+^ compared to HTkB and HMfB, respectively.

**Figure 2.**
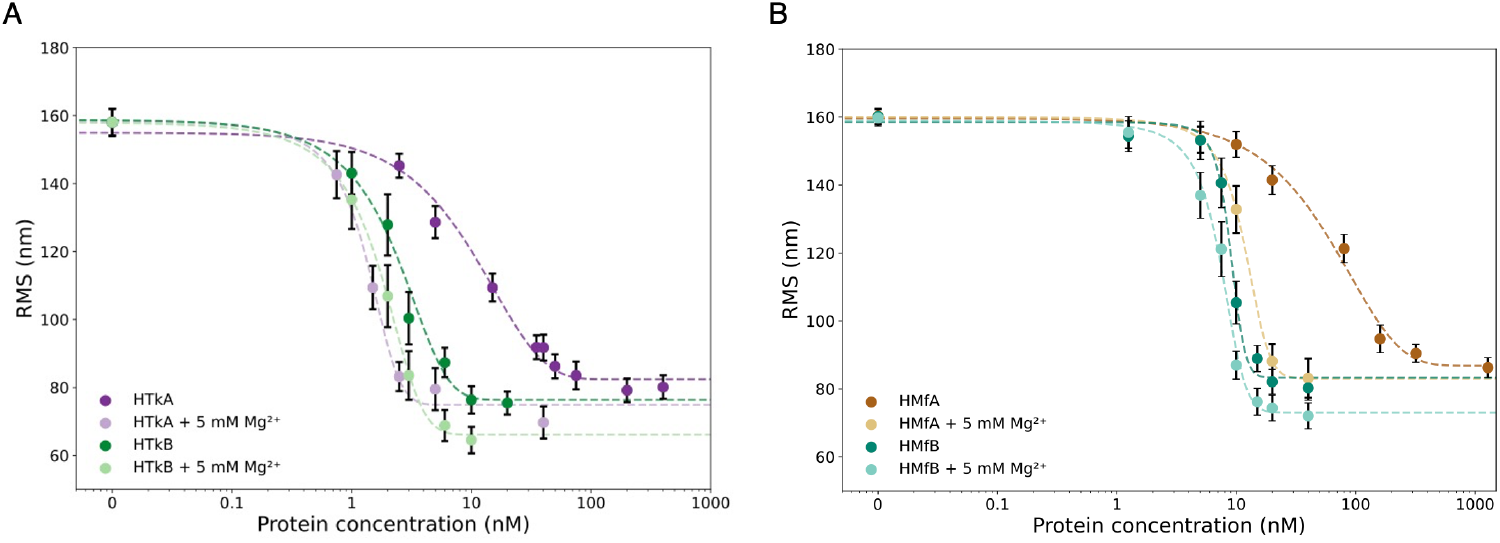
Mg^2+^ promotes formation and enhances HTkA/B and HMfA/B hypernucleosome compaction. A) In 5 mM MgC12,thc HTkA/B protein titration curves shift to lower concentrations, evidenced by the reduction in midpoint concentrations (see Table 1). The reduction in midpoint concentration indicates an effective increase in affinity. The increase in steepness suggests enhanced cooperativity. The lower RMS at saturating conditions implies a more tightly packed hypcmucleosome complex. (B) In 5 mM MgCl_2_, the HMfA/B protein titration curves show a similar trend in affinity, cooperativity, and compaction as HTkA/B. Each measurement (N > 100) was done in duplicate, with the error bars representing the standard deviation. Dashed lines are added to guide the eye.

For HTkA, the presence of 5 mM Mg^2+^ reduced the midpoint concentration from 9.2 ± 1.1 nM to 1.33 ± 1.1 nM and increased the steepness of RMS decrease, suggesting an increase in DNA affinity and cooperativity. At protein saturation, we measured an RMS of 69 ± 5 nm and an EED of 22 ± 11 nm in 5 mM Mg^2+^, compared to an RMS of 80 ± 4 nm and an EED of 33 ± 3 nm in absence of Mg^2+^, indicating the formation of more tightly packed structures. Experiments with HTkB also yielded a decrease in midpoint concentration from 2.5 ± 1.1 nM to 1.8 ± 1.1 nM. At protein saturation, an RMS of 64 ± 5 nm and an EED of 19 ± 9 nm in 5 mM Mg^2+^ was measured, compared to an RMS of 75 ± 4 nm and an EED of 26 ± 8 nm without Mg^2+^, implying an increased compaction. The HTkB_K27A,K62A,K66A_ stacking mutant demonstrated a similar trend with a decreased midpoint concentration and EED in Mg^2+^ (Supplementary Figure 2) Similarly, the presence of 5 mM Mg^2+^ lowered the midpoint concentration of HMfA from 59.7 ± 1.2 nM to 11.6 ± 1.1 nM. A shift in HMfB midpoint concentration was observed as well, from 9.0 ± 1.0 nM to 7.3 ± 1.1 nM in the presence of 5 mM Mg^2+^. Altogether, these results suggest that Mg^2+^ promotes the formation of hypernucleosomes, irrespective of the residues that play a crucial role in histone stacking, and enhances the compaction of the hypernucleosome complex.

Mg^2+^ titration experiments in TPM at constant protein concentration, chosen to allow observation of possible changes, confirmed the effect of Mg^2+^ on hypernucleosome formation (Figure 3). For 5 nM HTkA, a decrease in RMS from 140 nm to 100 nm was observed in the presence of 0.5 mM Mg^2+^ (Figure 3A). When the Mg^2+^ concentration was increased to 5 mM, the RMS further decreased to 80 nM (Figure 3A). For 0.5 nM HTkB, the RMS dropped from 130 nm to 110 nm in the presence of Mg^2+^ (Figure 3A). A comparable reduction was observed for HTkB_K27A,K62A,K66A_, for which the RMS decreased from 150 nm to 90 nm in the presence of 5 mM Mg^2+^ (Supplementary Figure 2). Similar titration in HMfA/B corroborated that Mg^2+^ promotes hypernucleosome formation (Supplementary Figure 3). To exclude the possibility that the observed decrease in RMS is due to an increase in ionic strength, we also performed titration experiments in HTkA, wherein the KCl concentration was adjusted to compensate for the increase in Mg^2+^ ions (Figure 3B). No significant difference was observed compared to the measurements without compensating for the ionic strength. Furthermore, increasing the K^+^ concentration in the absence of Mg^2+^ did not cause a decrease in RMS (Figure C). These results indicate that Mg^2+^ specifically affects hypernucleosome formation and compaction, and this cannot be simply attributed to a change in ionic strength.

**Figure 3.**
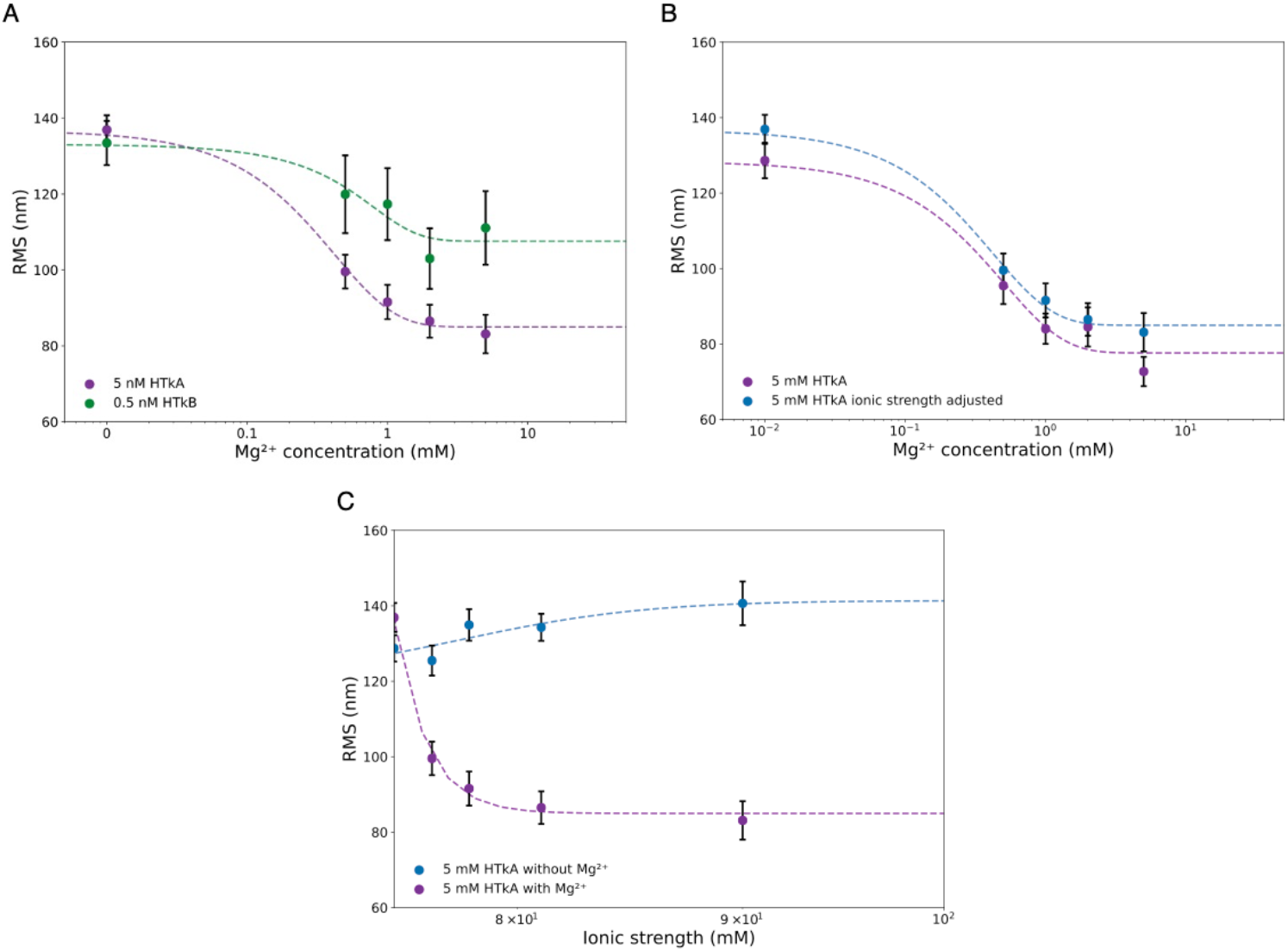
Mg^2+^ promotes archaeal hypernucleosome formation. **A)** Titration experiments with increasing MgCh at concentrations of 5 nM HTkA and 0.5 nM HTkB reveal an Mg^2+^ concentration-dependent increase of archaeal nucleosome formation. B) Mg^2+^ titration experiments with 5 nM HTkA in which the K^+^ concentration was reduced to adjust for the addition of Mg^2+^, maintaining a constant ionic strength throughout the Mg^2+^titration C) To further elucidate the effect of the ionic strength, a titration in 5 nM HTkA was conducted in the absence of Mg^2+^ while increasing K? concentration to achieve the same ionic strength (blue line). These results show that the compaction can be attributed directly to Mg^2+^ rather than to changes in ionic strength. Each measurement (N > 100) was performed in duplicates, with the error bars representing the error propagated standard deviation. Dashed lines are added to guide the eye.

### Single-molecule force spectroscopy resolves disparate Mg^2+^-mediated effects on HTk and HMf hypernucleosome stability

In previous studies, we used single-molecule force spectroscopy to measure the force required to unfold a hypernucleosome to the full contour length of the DNA in a two-step process wherein first, the interactions between stacking histone dimers rupture at several pN, followed by full unwrapping of the histones around 15 pN. A statistical mechanics model captured this behavior (28). For an extensive description of the model, see Supplementary Method. Fitting the force-distance curves to this model yields the number of dimers bound to DNA, stacking energy, wrapping energy, stiffness, and bending angle. The stacking energy reflects all interactions contributing to the stability of the stacked state. Here, we fit this statistical mechanics model to different archaeal histone-DNA complexes and investigate how these parameters depend on the Mg^2+^ concentration, providing a more detailed picture of how hypernucleosome stability is affected by Mg^2+^.

First, we characterized the mechanical properties of the HTkA and HTkB hypernucleosomes. In vitro reconstitution was done on a torsionally-unconstrained DNA substrate using 100 nM HTkA, and 20 nM HTkB, conditions representing saturation, and thus, complete coverage of DNA by histones. Figure 4A shows the force-distance curve of the HTkB hypernucleosome in the absence of Mg^2+^, exhibiting similar features as reported for HMfA/B fibers (28). Similar to HMfA/B, no hysteresis was observed between the stretch and release curves (Supplementary Figure 5) suggesting that the HTkB-DNA hypernucleosome complex is in thermal equilibrium during the stretching cycle. The fitted parameters are shown in Table 2. In the presence of 2 mM Mg^2+^, higher forces are required to perturb both the stacking and the wrapping of HTkB (Figure 4A).

**Table 2.**
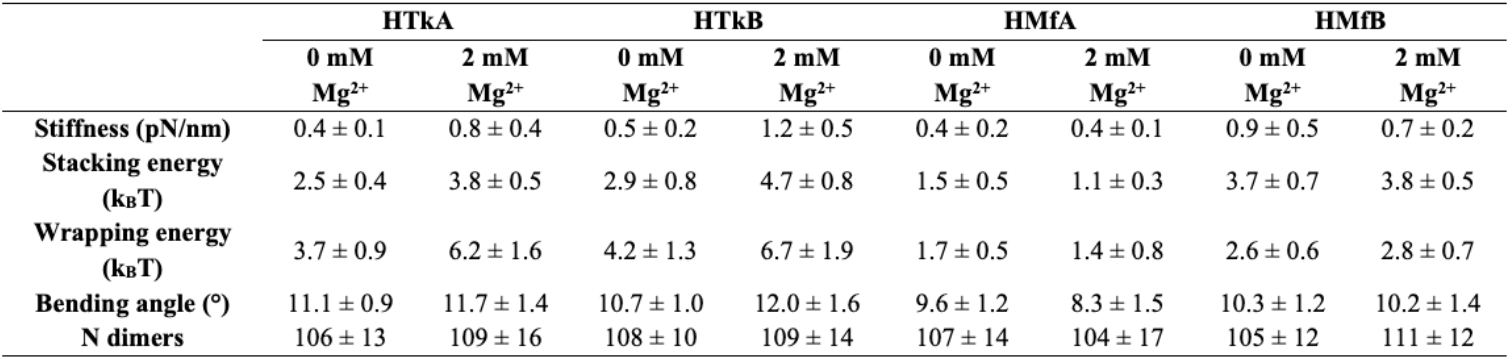
Fit of the statistical physics model (Supplementary Method) results to the force-extension curves of HTkA, HTkB, HMfA, and HMfB complexes in the presence and absence of 2 mM Mg^2+^.

**Figure 4.**
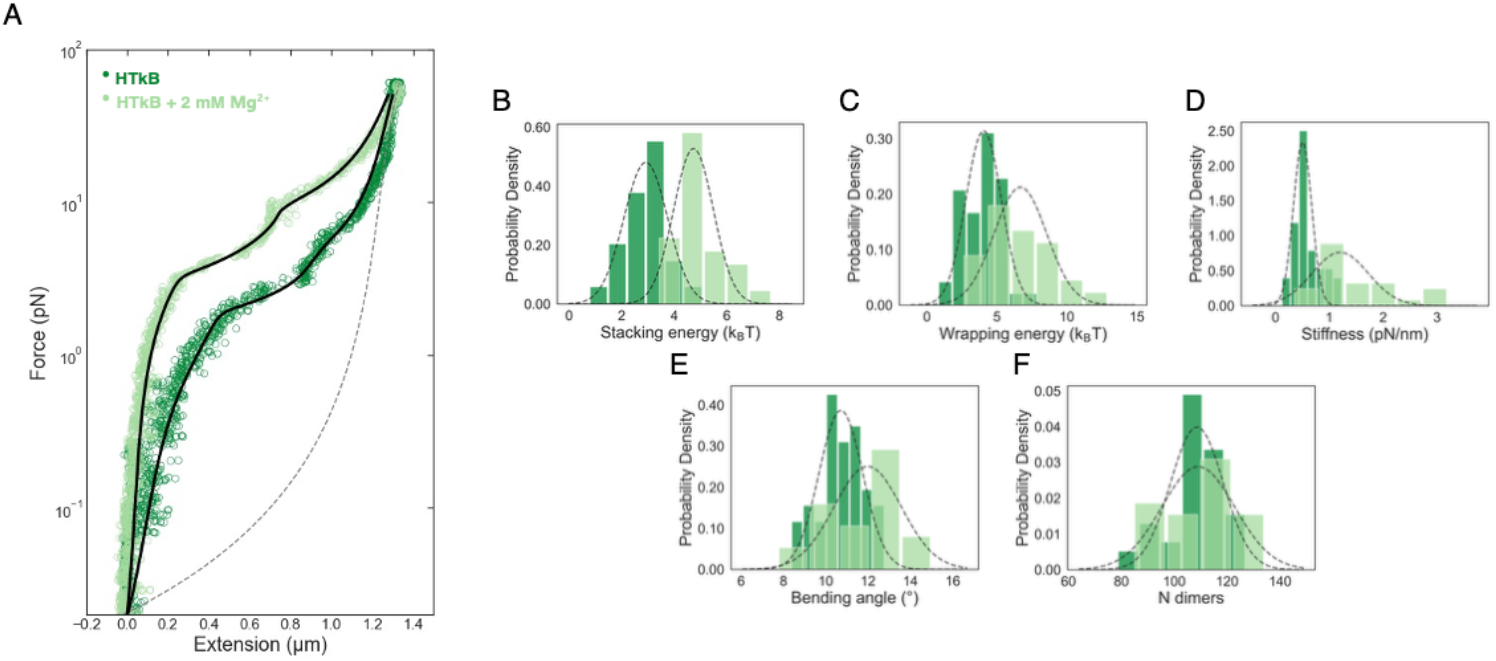
Mg^2+^ increases the compaction of HTkB fibers. **A)** Force extension curves and fits of HTkB hypcmuclcosomc in the absence (g_stack_=2.85 k_B_T, g_wrap_=3.67 kuT, stiffhcss=0.51 pN/nm and anglc=10.23°) and the presence of 2 mM Mg^2+^ions (g_stack_=4.24 k_B_T, g_wrap_=6.49 knT, stiffness=0.91 pN/nm and angle=11.61°), as indicated. The force required to reach extension of DNA, depicted by the dashed grey line, is higher in the presence of Mg^2+^ ions than in the absence of Mg^2+^ ions, representing a more stable hypemucleosomc complex. This can be directly attributed to the Mg^2+^ ions. B-F) The histograms of stacking energy, wrapping energy, stiffness, bending angle, and number of bound dimers of HTkB in the absence and the presence of Mg^2+^ ions. The histograms were fitted with a normal distribution. The mean and standard error of mean of all fit parameters can be found in Table 2.

For HTkA, we observe a similar response in the force-distance curve as for HTkB (Supplementary Figure 6 and Table 2). The previously observed tighter packing of the hypernucleosome is reflected by an increase in the stacking and wrapping energies as well as in hypernucleosome stiffness, confirming that Mg^2+^ stabilizes the HTkA and HTkB hypernucleosome complexes.

We then investigated if the stabilizing effect of Mg^2+^ is conserved across hypernucleosomes formed by other histones by performing the same series of single-molecule force spectroscopy experiments on HMfA and HMfB. Strikingly, we did not observe increased stacking and wrapping energies or stiffness of HMfB hypernucleosomes in the presence of Mg^2+^ (Figure 5 and Table 2), implying no increase in hypernucleosome complex stability. No change was observed in the presence of Mg^2+^ for HMfA either (Supplementary Figure 7 and Table 2). However, the stacking and wrapping energies for HMfB are significantly higher than for HMfA, in agreement with our previous study (28).

**Figure 5.**
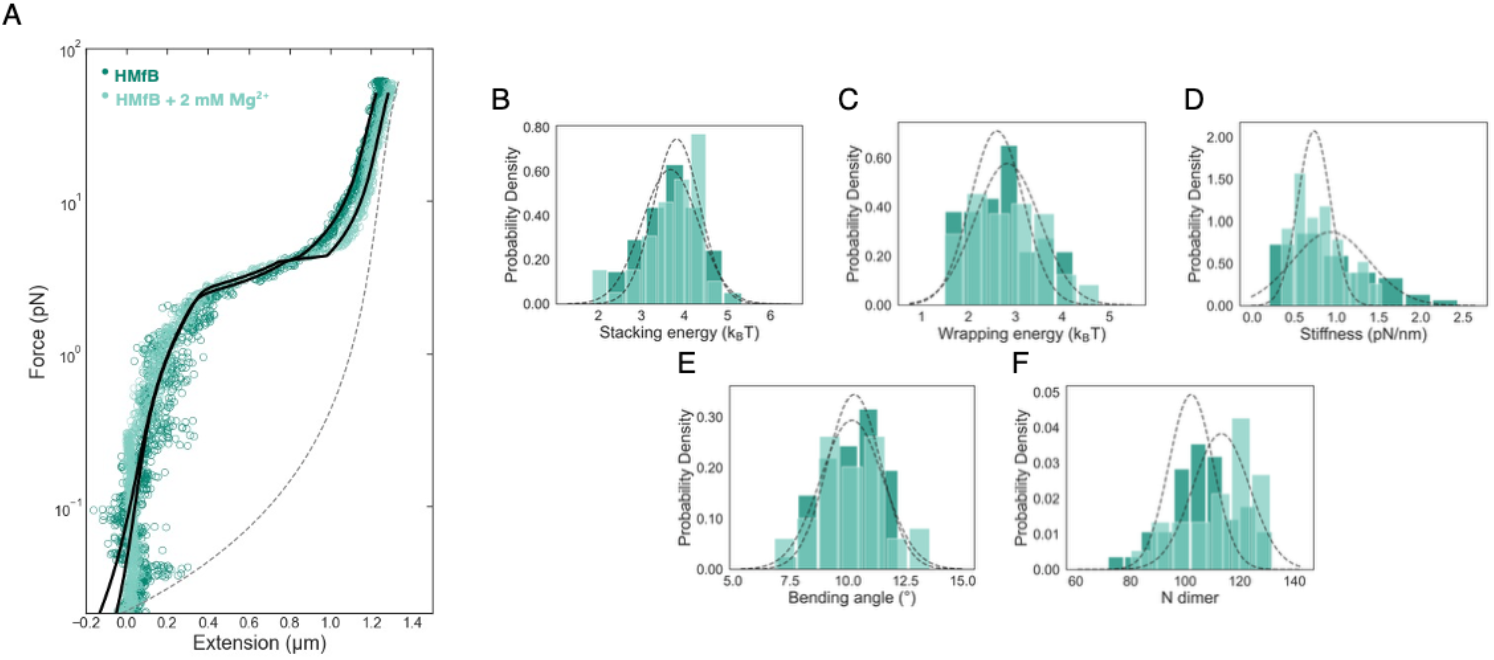
Mg^2+^ does not affect the compactness of HMfB fibers. **A)** Force-extension curves and fits of HMfB hypernucleosome in the absence (g_stack_=3.64 k_B_T, g_wrap_=2.78 k_B_T, stiffhess=0.81 pN/nm and angle=9.99°) and the presence of 2 mM Mg^2+^(g_slack_=3.99 k_B_T, g_wrap_=2.42 k_B_T, stiffhess=0.78 pN/nm and angle=10.2°), as indicated. Unlike the IITk histones, we do not observe a noticeable difference between the traces, also reflected by MST experiments. B-F) The histograms of stacking energy, wrapping energy, stiffness, bending angle, and number of bound dimers of HTkB in the absence and the presence of Mg^2+^. The histograms were fitted with a normal distribution. The mean and standard error of mean of all fit parameters can be found in Table 2.

**Figure 7.**
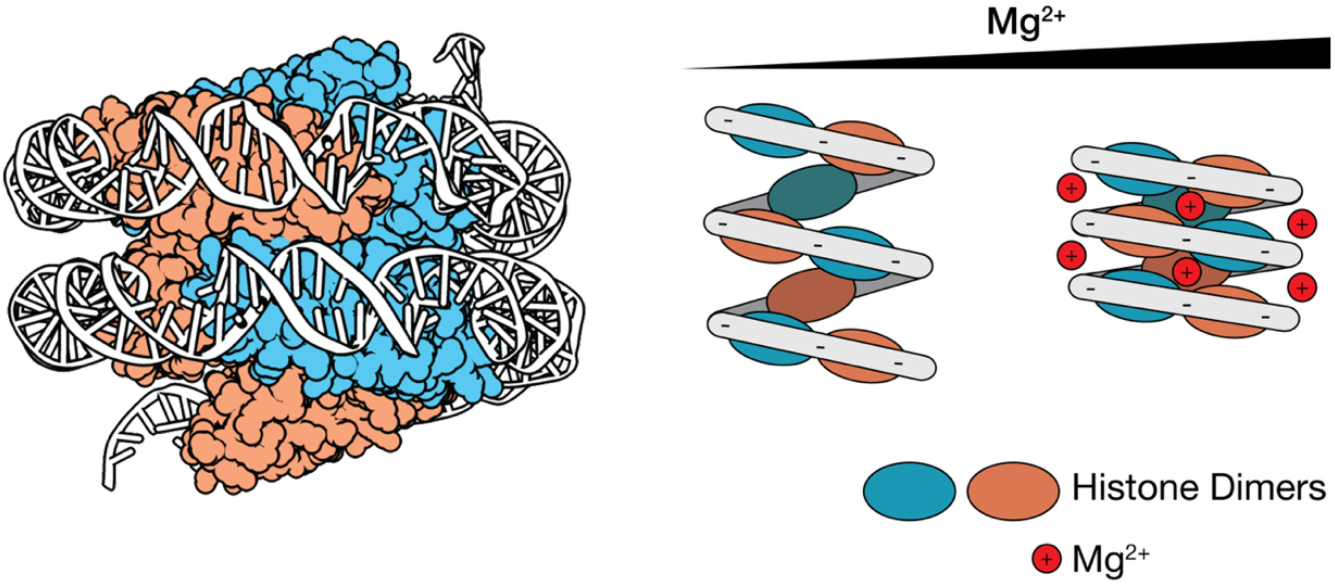
Modulation of archaeal hypernucleosome structure by Mg^2+^. (A) HMfB hypernucleosome structure based on data by Mattiroli et al. (2017) (PDB: 5T5K). (B) The presence of Mg^2+^ in solution shields the electrostatic repulsion of the negatively charged backbone, promoting a more compact hypernucleosome conformation.

## DISCUSSION

The structural and mechanical properties of archaeal hypernucleosomes have been thoroughly characterized during the last decade. At least as important, however, is determining how this structure is modulated in the context of genome accessibility. Through this study, we characterized in solution how environmental conditions may play a role in this modulation. We first demonstrated that hypernucleosome formation is a conserved feature of canonical archaeal histone variants. Furthermore, we showed that Mg^2+^ universally promotes the formation and compaction of hypernucleosomes, and in the case of HTkA/B hypernucleosomes, increases hypernucleosome stability. These findings may be relevant to how hypernucleosome accessibility in archaea is regulated. Most archaeal histones lack the N-terminal tails, as compared to the long N-terminal tails of histones in eukaryotic nucleosomes which are typically functionally modulated via post-translational modification. One of Nature’s solutions may be the post-translational modification of residues at the hypernucleosomal stacking interface (30). However, sensitivity of histones to environmental cues, such as changes in the Mg^2+^ concentration, may be an alternative mechanism to regulate genome accessibility.

Our TPM experiments reveal that HTkA and HTkB assemble into hypernucleosomes on long DNA substrates, similar to earlier findings regarding HMfA/B histones (28). This also corroborates previous *in vitro* and *in vivo* observations (26,38,41). Of note, both *M. fervidus* and *T. kodakarensis* utilize a pair of histone paralogs that, by having distinct individual properties, may function in providing a means of controlling chromatin structure, either directly through differential DNA binding, or indirectly, through possible heterodimerization. *In vivo*, the differential expression of HTkA (1.1% of the proteome) and HTkB (0.66%) during the exponential growth phase (11) indeed supports a model of chromatin regulation mediated by an interplay between histone paralogs. Remarkably, the histone paralogs exhibit a differential response to Mg^2+^, as well. Both HMfA and HTkA are more sensitive to changes in Mg^2+^ than HMfB and HTkB respectively. It can be noted that at position 14, which is predicted to facilitate stacking interaction, an asparagine is present in HMfA as opposed to aspartic acid in HMfB. At position 66, which is predicted to form the tetramerization interface, a methionine is present in HMfA, in contrast to arginine in HMfB. (Supplementary Figure 1). Both of these differences imply that salt bridges that stabilize both dimerization and tetramerization are reduced in HMfA compared to HMfB. Overall, this may cause the formation of a less compact and stable hypernucleosome, explaining why Mg^2+^-induced compaction is enhanced in HMfA hypernucleosomes compared to HMfB. In relation to the *T. kodakarensis* histones, *in vitro* experiments have shown that HTkB has a higher binding affinity for DNA than HTkA (42). *In silico* analysis has revealed the conservation of residues unique to HTkA-like histones (E18, Y35) and HTkB-like histones (K35). Molecular dynamics have confirmed the higher DNA binding affinity for HTkB compared to HTkA (11). However, the residues predicted to be involved in stacking interactions do not differ between HTkB and HTkA (K14, K26, E30, E34, Q48, E58, K61, K65) (10), making it unclear what molecular difference leads to the formation of a more stable hypernucleosome with HTkB, and why HTkA is more sensitive to changes in Mg^2+^.

While the TPM experiments revealed enhanced compaction of DNA in HMfA/B hypernucleosomes, the presence of Mg^2+^ did not induce any changes in cooperativity or affinity, which was confirmed in the magnetic tweezer experiments. This suggests that the changes induced by Mg^2+^ measured in the TPM experiments should be attributed to electrostatic shielding of the DNA, which is known to be a significant factor in compaction of eukaryotic nucleosomes (34,36). This shielding, in turn, allows for the DNA gyres within the hypernucleosome to be closer. Indeed, conducting the same experiments on HTkB_K27A,K62A,K66A_, which lack the residues necessary for stacking showed that enhanced compaction of the hypernucleosome in the presence of Mg^2+^ is maintained, suggesting that the electrostatic shielding is a universal effect.

Both *T. kodakarensis* and *M. fervidus* are extremophiles, as shown by their optimal growth temperatures and the environmental conditions in which they thrive. In the natural habitat of *T. kodakarensis*, high levels of Mg^2+^, Fe^2+/3+^, Mn^2+^, and Cd^2+^ were found (43) and the intracellular concentration of Mg^2+^ in *T. kodakarensis* was estimated to be around 120 mM (44), much higher than in eukaryotes. Therefore, *T. kodakarensis* likely has adapted mechanisms to cope with varying Mg^2+^ concentrations, possibly affecting genome accessibility through histone stacking and consequently influencing gene expression in response to changes in Mg^2+^ concentrations. In contrast, *M. fervidus* thrives in Mg^2+^-deficient environments (45) and may therefore be less affected by the presence of Mg^2+^. This difference in ecological niche could account for the varying effects of Mg^2+^ on hypernucleosome stability between the two organisms.

The interplay between hypernucleosome formation, compaction, and stabilization, its role in gene regulation, and its dependence on the presence of cations presents a regulatory mechanism whose precise details need further study. Our current findings highlight the role of Mg^2+^ in hypernucleosome assembly and compaction. Furthermore, we demonstrate that this effect is not equally strong for HTk and HMf, despite their high structural similarity. With the knowledge of diverse variants of archaeal histones and the potential interplay among them, gaining a better understanding of the molecular mechanisms underlying this distinction is essential for obtaining further insights into the function of structural regulation of archaeal hypernucleosomes in archaeal chromatin and their role in transcription regulation in response to environmental cues.

## MATERIALS AND METHODS

### Recombinant histone purification

HMfA and HMfB were purified as described (Henneman et al., 2021). The purity and integrity of proteins was verified through intact protein liquid chromatography/mass spectroscopy (LC/MS). Accurate protein folding was confirmed with circular dichroism (CD) spectroscopy. The native sequence of genes *htkA* (TK1413) and *htkB* (TK2289) was synthesized (GeneArt/ThermoScientific) and cloned into pET30b via Gibson assembly (Gibson et al., 2009). The sequence was verified through Sanger sequencing. The plasmids for expression of HTkA (pRD457) and HTkB (pRD459) are available from Addgene. Following transformation into Rosetta (DE3) pLySs cells (Novagen), the cells were grown in LB with 50 μg/mL kanamycin and 25 μg/mL chloramphenicol at 37 °C, 220 rpm to an OD_600_ of 0.4. Protein expression was induced with 1 mM IPTG; the culture was further incubated for 3 hours at 37 °C, 220 rpm. Cells were harvested at 7,510 xg for 30 minutes at 4 °C, resuspended in 50 mM NaCl, 25 mM Tris, pH 7.0, and lysed using a Stansted Pressure Cell Homogenizer S-PCH-10 at 310 MPa. The lysate was centrifuged at 4 °C for 20 minutes at 10,000 rpm (Beckmann Coulter, type 70Ti Fixed-angle Titanium rotor). The supernatant was subjected to 40 μg/mL DNase I for 1 hour at 37 °C (after adding MgCl_2_ up to a concentration of 5 mM) to digest DNA. Next, it was incubated at 75 °C for 20 minutes to denature endogenous *E. coli* proteins while retaining the thermostable heterologously expressed histones. Centrifugation was repeated for 30 minutes at 10,000 rpm (Beckmann Coulter, type 70Ti Fixed-angle Titanium rotor), and the supernatant was filtered and loaded onto a 5 mL HiTrap Heparin HP column (Cytiva) using a salt gradient (50 mM to 1 M of NaCl in 25 mM Tris-HCl, pH 7.0). SDS-PAGE was used to identify fractions containing histones. Next, these fractions were pooled, concentrated and subjected to size exclusion chromatography on a Superdex 75 Increase 10/300 GL column (Cytiva) using 600 mM NaCl in 25 mM Tris-HCl, pH 7, as buffer. Fractions were again analyzed using SDS-PAGE. The histone-containing fractions were pooled, dialyzed overnight against 100 mM KCl, 25 mM Tris-HCl, pH 7.0., and protein concentrations were determined with a Qubit 4 fluorometer. The purity and integrity of the protein were verified through intact protein LC/MS. Accurate protein folding was confirmed with CD spectroscopy.

### Tethered particle motion

The tethered particle motion (TPM) experiments were performed and analyzed as described (Henneman et al., 2024; Hu et al., 2024). Using a 685 bp nonspecific DNA substrate in 25 mM Tris-HCl, pH 7.0, 75 mM KCl, and 5 mM MgCl_2_, each measurement was done in duplicate. To select for single-tethered beads, an anisotropic ratio cut-off of 1.3 and a standard deviation cut-off of 8% was set. Means and standard deviations of the individual measurement series for each concentration were calculated by maximum likelihood estimation assuming a normal distribution. Outliers with a Z-score >3 or <-3 were not considered for fitting. The “line to guide the eye” was generated by fitting the means to a logistic function 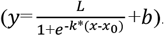. A custom Python script was used for fitting and plotting the TPM data. For plotting, the means of the two individual measurements were averaged for each measured concentration and the standard deviations were error-propagated 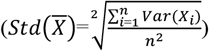. The end-to-end distance (EED) was calculated from the 5% largest deflection of all beads in the XY-plane by subtracting the mean bead radius, as described in Henneman et al. (2021). The calculated values were then fitted to a sigmoidal function, and from the fit we obtained the error value and error (Supplementary Figure 3).

### Magnetic tweezers

Magnetic tweezer experiments were done according to an established protocol (Brouwer et al., 2024; Henneman et al., 2021) in 25 mM Tris-HCl, pH 7, 75 mM KCl, and 2 mM MgCl_2_, as necessary. A force ramp of 0 to 60 pN was applied in 70 seconds, with a 1-second hold before the force was released at the same rate. This was done twice, with a 1-second pause in between each cycle. Analysis was done by fitting a previously devised statistical physics model to the data (Henneman et al., 2021) (Supplementary Method).

## DATA AVAILABILITY

Raw data are available from the 4TU repository (https://data.4tu.nl) with the DOI: 10.4121/bda6e3ba-cd39-4db8-9568-87e4d503c59f

## AUTHOR CONTRIBUTIONS

Ilias Zarguit: Conceptualization, Data curation, Formal analysis, Investigation Methodology, Validation, Visualization, Writing—original draft. Marc Kenneth Cajili: Conceptualization, Data curation, Formal analysis, Investigation Methodology, Validation, Visualization, Writing—original draft Bert van Erp: Methodology, Investigation. Samuel Schwab: Software. Nico van der Vis: Investigation, Methodology. Marianne Bakker: Investigation, Methodology. John van Noort: Software, Methodology, Writing—review & editing, Resources, Supervision. Remus T. Dame: Conceptualization, Methodology, Writing—review & editing, funding acquisition, Project administrator, Resources, Supervision.

## ACKNOWLEDGEMENTS

We would like to thank Antonella Di Savino for help with the purification of HTkA and HTkB.

