## Supplementary information for "MODULATION OF ARCHAEAL HYPERNUCLEOSOME STRUCTURE AND STABILITY BY Mg^2+^"

**Supplementary Table 1. Oligonucleotides used for this study**

| Primer Name | Plasmid | Sequence (5′ – 3′) |
|---|---|---|
| HTkA insert F | pRD457 | GAAATAATTTTGTTTAACTTTAAGAAGGAGATATACATATGGCAGAACTGCCGATTGC |
| HTkA insert R | pRD457 | CCTTTCGGGCTTTGTTAGCAGTTAGGCCTTGATGGCCAGTTTAATATCTTC |
| HTkA vector F | pRD457 | GAAGATATTAAACTGGCCATCAAGGCCTAACTGCTAACAAAGCCCGAAAGG |
| HTkA vector R | pRD457 | GCAATCGGCAGTTCTGCCATATGTATATCTCCTTCTTAAAGTTAAACAAAATTATTTC |
| HTkB insert F | pRD459 | GAAATAATTTTGTTTAACTTTAAGAAGGAGATATACATATGGCAGAACTGCCGATTGC |
| HTkB insert R | pRD459 | CCTTTCGGGCTTTGTTAGCAGTTAGCTTTTGATGGCCAGTTTAATATCTTCTGC |
| HTkB vector F | pRD459 | GCAGAAGATATTAAACTGGCCATCAAAAGCTAACTGCTAACAAAGCCCGAAAGG |
| HTkB vector R | pRD459 | GCAATCGGCAGTTCTGCCATATGTATATCTCCTTCTTAAAGTTAAACAAAATTATTTC |

### Substrates used for the study

### TPM substrate

5’TTACTTTCACCAGCGTTTCTGGGTGAGCAAAAACAGGAAGGCAAAATGCCGCAAAAAAGGGAATAAGGGCGAC
ACGGAAATGTTGAATACTCATACTCTTCCTTTTTCAATATTATTGAAGCATTTATCAGGGTTATTGTCTCATGAG
CGGATACATATTTGAATGTATTTAGAAAAATAAACAAATAGGGGTTCCGCGCACATTTCCCCGAAAAGTGCCACC
TGACGTCTAAGAAACCATTATTATCATGACATTAACCTATAAAAATAGGCGTATCACGAGGCCCTTTCGTCTTCA
AGAATTCCGGCGCAAATTCGTGACCAGTTGCATCAGCTGCGTGAGCTGTTTATCGCAGCATCGTAACAGGATAGT
GAAGAAGACTAAGCTTTAATGCGGTAGTTTATCACAGTTAAATTGCTAACGCAGTCAGGCACCGTGTATGAAATC
TAACAATGCGCTCATCGTCATCCTCGGCACCGTCACCCTGGATGCTGTAGGCATAGGCTTGGTTATGCCGGTACT
GCCGGGCCTCTTGCGGGATATCGTCCATTCCGACAGCATCGCCAGTCACTATGGCGTGCTGCTAGCGCTATATGC
GTTGATGCAATTTCTATGCGCACCCGTTCTCGGAGCACTGTCCGACCGCTTTGGCCGCCGCCCAGTCCTGCTCGC
TTCGCTACTTGG 3’

### MT substrate

5’ACCGCGAGACCCACGCTCACCGGCTCCAGATTTATCAGCAATAAACCAGCCAGCCGGAAGGGCCGAGCGCAGA
AGTGGTCCTGCAACTTTATCCGCCTCCATCCAGTCTATTAATTGTTGCCGGGAAGCTAGAGTAAGTAGTTCGCCA
GTTAATAGTTTGCGCAACGTTGTTGCCATTGCTACAGGCATCGTGGTGTCACGCTCGTCGTTTGGTATGGCTTCA
TTCAGCTCCGGTTCCCAACGATCAAGGCGAGTTACATGATCCCCCATGTTGTGCAAAAAAGCGGTTAGCTCCTTC
GGTCCTCCGATCGTTGTCAGAAGTAAGTTGGCCGCAGTGTTATCACTCATGGTTATGGCAGCACTGCATAATTCT
CTTACTGTCATGCCATCCGTAAGATGCTTTTCTGTGACTGGTGAGTACTCAACCAAGTCATTCTGAGAATAGTGT
ATGCGGCGACCGAGTTGCTCTTGCCCGGCGTCAATACGGGATAATACCGCGCCACATAGCAGAACTTTAAAAGTG
CTCATCATTGGAAAACGTTCTTCGGGGCGAAAACTCTCAAGGATCTTACCGCTGTTGAGATCCAGTTCGATGTAA
CCCACTCGTGCACCCAACTGATCTTCAGCATCTTTTACTTTCACCAGCGTTTCTGGGTGAGCAAAAACAGGAAGG
CAAAATGCCGCAAAAAAGGGAATAAGGGCGACACGGAAATGTTGAATACTCATACTCTTCCTTTTTCAATATTAT
TGAAGCATTTATCAGGGTTATTGTCTCATGAGCGGATACATATTTGAATGTATTTAGAAAAATAAACAAATAGGG
GTTCCGCGCACATTTCCCCGAAAAGTGCCACCTGACGTCTAAGAAACCATTATTATCATGACATTAACCTATAAA
AATAGGCGTATCACGAGGCCCTTTCGTCTCGCGCGTTTCGGTGATGACGGTGAAAACCTCTGACACATGCAGCTC
CCGGAGACGGTCACAGCTTGTCTGTAAGCGGATGCCGGGAGCAGACAAGCCCGTCAGGGCGCGTCAGCGGGTGTT
GGCGGGTGTCGGGGCTGGCTTAACTATGCGGCATCAGAGCAGATTGTACTGAGAGTGCACCATATGCGGTGTGAA
ATACCGCACAGATGCGTAAGGAGAAAATACCGCATCAGGCGCCATTCGCCATTCAGGCTGCGCAACTGTTGGGAA
GGGCGATCGGTGCGGGCCTCTTCGCTATTACGCCAGCTGGCGAAAGGGGGATGTGCTGCAAGGCGATTAAGTTGG
GTAACGCCAGGGTTTTCCCAGTCACGACGTTGTAAAACGACGGCCAGTGCCAAGCTTGCATGCCTGCAGGTCGAC
GAGGTGATATTTCCAATTTGGGAAATTTCCCAAATCAGTAATGTAGCCTCTACGGGTGTCTCTGTCAGCCCCGTG
GTCGCCAGCACAGAATGTATCGTACCCCTGAAGGTAGTTTTTACCGCCGTGGCACACGATAAAGGTGCACCTTG
TGATAATAAGGTGGAAAAATATATATGAAAAAGTGAAATTGATTGTGGCTGCACTAGGACATCATTATTTCTTAC
TTGGCTATTTACACGTACTTACGCTGGCTGTATATCATTTAAGGGGCGGAGGACGAAGAGGACGGACCCGAGATC
ATCCGGTCCAAGAAACGGGTCATCCGGTCCTTAGCATTGTCTAGACTATCTAGGGCAGGACGGACATCCACGTGG
AAAGTAGGCATTCCGTTTTCGTCGTCGGGCCCTCCGTAGAAATCCAAGACGTATCTAACTTCCTTGAAGGTTGGA
GGTTGTTGTTCCGCTTTGCGCTCGCCTCGGAGTACAATCCAGTCGTGCCTGTCGAATGGTAGTTCTTGGCTAAAA
TGGGACGGAAACAGTAGGCCGCACAGGTGCATCCAGCGAGCACGAGGGCTCAATACGCCCGGTTTCCCCATGAAT
TTCAGCAACTTAGGCTGCACGTGGCTTTCATCTGTGTGCGGTTTTTCCCATTCGAGCACTTCCTGCCAGCACCCT
TCATTTAGAAAGTTGTGGACCTGCACCATCGACTCCACTGCATCTTCGGCGACTTCGCCGCTACCGCCAATCTTG
CCCTTTCTAACCATAGCATTGTACATCTGTTGTGGAGAAGGATACTCCCAGAACTCGTTACTGTCTGGACTCTTG
GGGATGCTGGAGATGGTCCGATCAACGGGCAAGTCCATCTTTTGGCCAGGCTGTTTGGATGCTGCCAACTCCGGC
ATATTGTTCAGCGGGTTTATTCTATCGTTATCTCCCTGCATAACGGGGCACTCAGAGGATGGTGGCGACGACGAC
GACGACTCGTGCATGACTGGGCACCCTGACATGGATGATACTGCTGCCCCACCAATATCTTTGCCCGTAGTTTTT
TGATCTGCCCAAAACCAACCCATTTTTTGTGAATTCATAGTATCGATACCCTCCTAAGTTTTGTAATCTATAAAG
TTAGCAATTTAACTAAGTGTAAAAACTTAGCTAGCTTACTAAAAAGATAATTACTGAAAAGCTAGCTTGGCCGGA
TCCGATCGACAAAGGAAAAGGGGCCTGTTTACTCACAAGCTTTTTTCAAGTAGGTAATTAAGTCGTTTCTGTCTT
TTTCCTTCTTCAACCCACCAAACGCGTACTTGGTACCAGGAATATATTTCTTTGGGTTAGTCAAGTACTCTGACA
TGTTATTTTCGTCCCACAACACGTTTTTCTTGATATTGGCATCTGTGTACGAATACCCTTCAGCTTGACCAGAGT
GTCTGCCAAAGATACCATGCAAGTTTGGACCAACCTTATGTGGGCCACCCTTTTCCACGGTGTGGCATTGTAGAC
ATCTAGTCTTGAAAAGTGTAGCACCTTTCTTAGCAGAACCGGCCTTGAATTCAGCCATGGATATATCTCCTTCTT
AAAGTTAAACAAAATTATTTCTAGATTAGTTAGTTATGGATCCCCGGGTACCGAGCTCGAATTCGTAATCATGGT
CATAGCTGTTTCCTGTGTGAAATTGTTATCCGCTCACAATTCCACACAACATACGAGCCGGAAGCATAAAGTGTA
AAGCCTGGGGTGCCTAATGAGTGAGCTAACTCACATTAATTGCGTTGCGCTCACTGCCCGCTTTCCAGTCGGGAA
ACCTGTCGTGCCAGCTGCATTAATGAATCGGCCAACGCGCGGGGAGAGGCGGTTTGCGTATTGGGCGCTCTTCCG
CTTCCTCGCTCACTGACTCGCTGCGCTCGGTCGTTCGGCTGCGGCGAGCGGTATCAGCTCACTCAAAGGCGGTAA
TACGGTTATCCACAGAATCAGGGGATAACGCAGGAAAGAACATGTGAGCAAAAGGCCAGCAAAAGGCCAGGAACC
GTAAAAAGGCCGCGTTGCTGGCGTTTTTCCATAGGCTCCGCCCCCCTGACGAGCATCACAAAAATCGACGCTCAA
GTCAGAGGTGGCGAAACCCGACAGGACTATAAAGATACCAGGCGTTTCCCCCTGGAAGCTCCCTCGTGCGCTCTC
CTGTTCCGACCCTGCCGCTTACCGGATACCTGTCCGCCTTTCTCCCTTCGGGAAGCGTGGCGCTTTCTCATAGCT
CACGCTGTAGGTATCTCAGTTCGGTGTAGGTCGTTCGCTCCAAGCTGG 3’

**Supplementary Method**

**Quantitative analysis of Magnetic tweezer data**

Henneman et al. (2021) devised a statistical physics model to describe the response of a hypernucleosome to force. The model takes into account three force dependent conformations of the hypernucleosome. In the first conformation, occurring at low force, the hypernucleosome is fully stacked, and all DNA is wrapped around the stacks of histones. In the low force regime the hypernucleosome behaves as a Hookean spring which is modeled by a freely jointed chain (FJC) as described by Equation 1.

$$Z_{FJC}(f)=L_{dimer}\left(coth\left(\frac{fb}{k_BT}\right)-\frac{k_BT}{fb}\right) \quad (1)$$

where $Z_{FJC}$ is the extension per dimer, $L_{dimer}$ represents the height of a single dimer, $f$ the applied force, $b$ the Kuhn length, $k_B$ Boltzmann's constant and T the temperature. In the FJC, this stiffness that is inversely proportional to the effective Kuhn length. At high forces, the FJC model converges to an asymptote, but at these forces the hypernucleosome ruptures into another conformation, as described below, so in practice Eq. 1 is only applied in the linear range.

In the second conformation, at intermediate forces, the DNA is still wrapped around histone dimers, but the histone dimers do not directly interact with each other. We use an extensible worm-like chain (eWLC) to model this regime (Equation 2).

$$Z_{WLC}(f)=L\left(1-\frac{1}{2}\sqrt{\frac{k_BT}{fP}}+\frac{f}{S}\right) \quad (2)$$

in which $Z_{WLC}$ is the extension of the histone-DNA dimer complex, L the contour length of the DNA in this complex, S the stretch modulus of DNA, f the force, T the temperature and P the persistence length of DNA. The eWLC chain is commonly used to describe the behavior of bare DNA. However, in case of DNA bound proteins, the trajectory of the DNA may include kinks. In that case the eWLC chain model still holds, but the persistence length reduces, depending on the bends induced by dimers (Equation 3).

$$P_{app}=\frac{P}{1+8N^2P\left(1-\frac{cos\frac{\alpha}{4}}{L}\right)^2} \quad (3)$$

where $P_{app}$ is the apparent persistence length, $N$ is the number of induced bends, and $\alpha$ is the protein-induced deflection angle.

At relatively high forces part of the histone-bound DNA unwraps. In this conformation the histones are still bound to the DNA, but do not deform it, resulting in a straight conformation where DNA is following the eWLC model as described before in Eq. 2., with a persistence length that equals that of bare DNA. The force-dependent free energy of each conformation can be calculated by:

$$G_{FJC}= \int z_{FJC}(f)df\text{-}g_{\text{stack}}\text{-}g_{wrap} \tag{4}$$

$$=L_{dimer}\frac{k_bT}{b}\left(ln\left(sinh\frac{fb}{k_bT}\text{-}ln\left(\frac{fb}{k_bT}\right)\right)\right)\text{-}g_{\text{stack}}\text{-}g_{wrap}\ )$$

$$G_{WLC}= \int z_{WLC}(f)df\text{-}g_{wrap}=L\left(f\text{-}\sqrt{\frac{f^{*}k_bT}{P}}+\frac{f^2}{2S}\right)\text{-}g_{wrap} \tag{5}$$

Here $g_{\text{stack}}$ and $g_{\text{wrap}}$ represent the protein-protein interaction energy and protein-DNA interaction energy respectively. In case of the straight conformational state, Eq. 5 can still be used, provided that $g_{\text{wrap}}$ is equal to 0.

The total free energy of the chromatin fiber in the magnetic tweezers is then obtained by summing the free energy contributions of each histone dimer in the hypernucleosome and subtracting the work performed by the bead *W(f)*:

$$G_{total}=G_{FJC}+G_{WLC}\text{-}W(f) \tag{6}$$

The extension of the HMf-DNA complex was calculated as the Boltzmann-weighted mean extension over all conformations *j*:

$$\langle z_{total}(f)\rangle = \frac{\sum_j z_j(f)e^{\frac{-G_j(f)}{k_BT}}}{\sum_j e^{\frac{-G_j(f)}{k_BT}}}\text{-}z_0 \tag{7}$$

in which an arbitrary offset $z_0$ accounts for the precise location of the DNA attachment to the bead.

Fitting Equation 7 to the magnetic tweezers data we are able to quantify the number of bound dimers, the stacking energy, the deflection angle, the stiffness of the hypernucleosome and the wrapping energy.

### Supplementary Figures

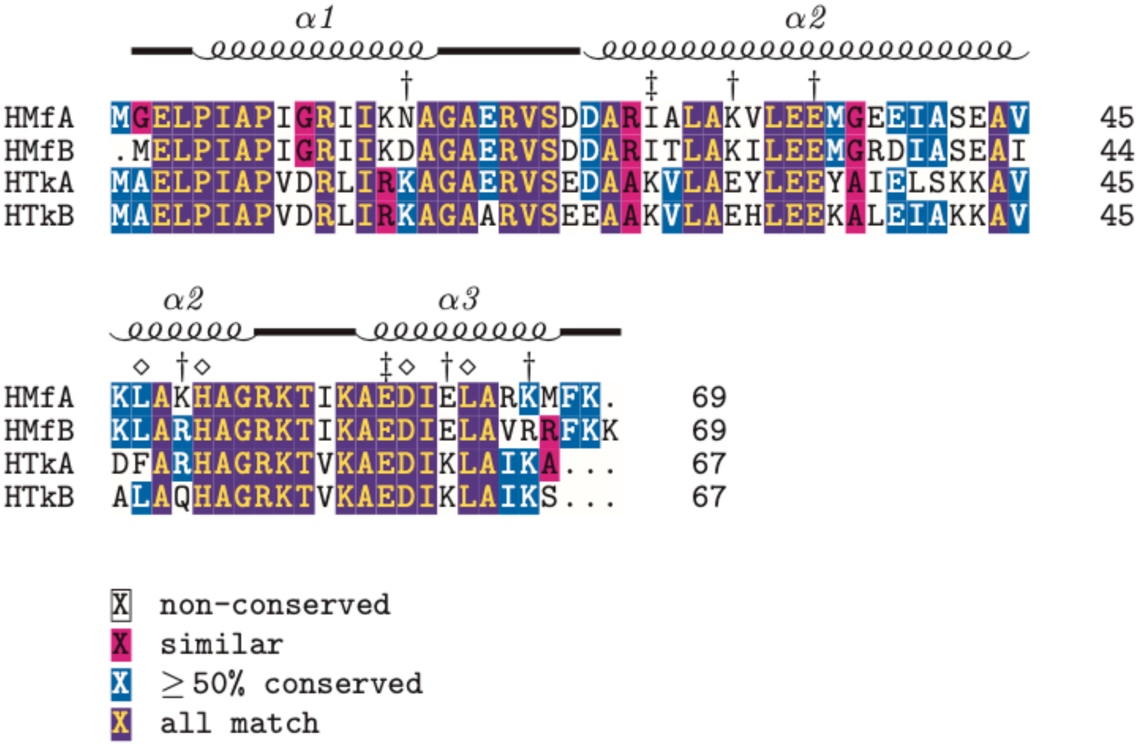


**Supplementary Figure 1.** Multiple sequence alignment of HMfA, HMfB, HTkA, and HTkB. Daggers (†) mark the predicted stacking residues for all histones (K30-E61, E34-R65, D14-R48 for HMf; E30-K61, E34-R65, K14-Q48 for HTk), while double daggers (‡) indicate stacking residues present in HTkA and HTkB but not HMfA and HMfB (K26-E58). Diamonds (◇) mark residues at the dimer-dimer interface.

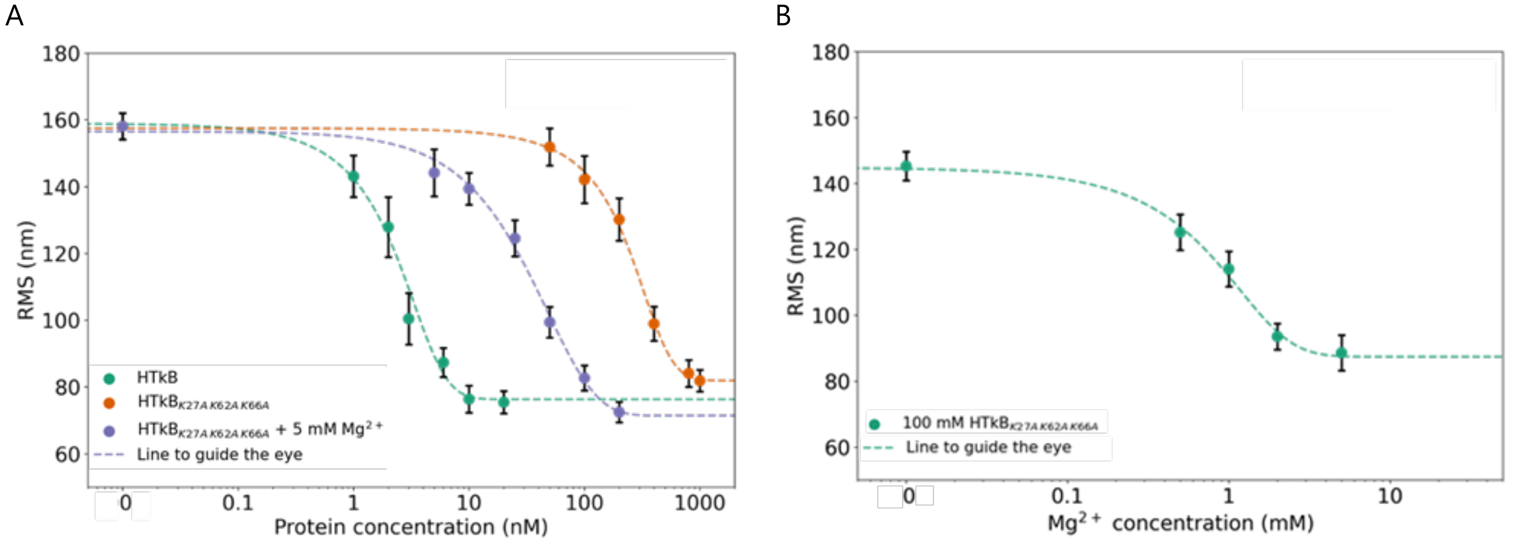


**Supplementary Figure 2. Tethered particle motion experiments reveal that hypernucleosomes with lysine-alanine substitutions that cancel stacking interactions of HTkB still form hypernucleosomes and that compaction is enhanced by $Mg^{2+}$.** (A) Tethered particle motion experiments demonstrate that the HTkB$_{K27A,K62A,K66A}$ stacking mutant can still form hypernucleosome structures on DNA substrates. Upon mutating the lysines involved in stacking into alanines, the midpoint concentration shifts from 2.46 ± 1.1 to 271.78 ± 1.1 nM, demonstrating that the affinity and the cooperativity are highly affected by the mutation. In the presence of 5 mM of $Mg^{2+}$ ions, the midpoint concentration shifts to 40.73 ± 1.2 nM, demonstrating that $Mg^{2+}$ ions promote the formation of hypernucleosome complexes also in absence of stacking interactions. B) $Mg^{2+}$ titration experiments show a decrease in RMS upon increased concentration of $Mg^{2+}$ ions.

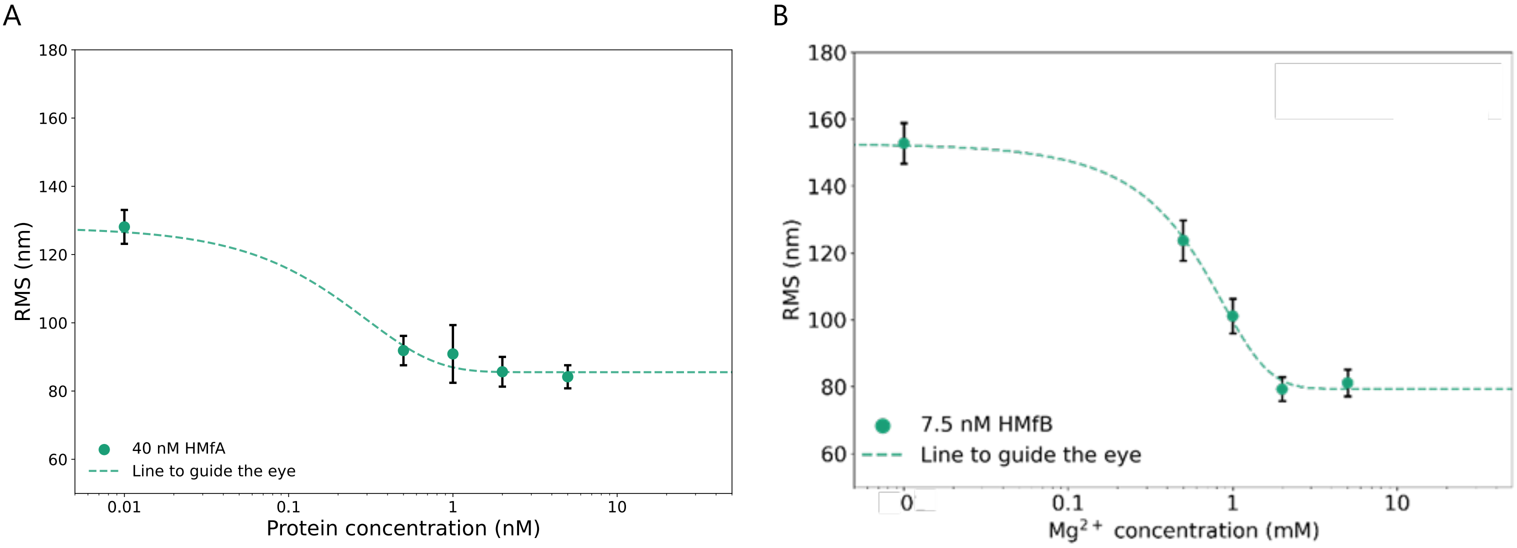


**Supplementary Figure 3.** Titration experiments with increasing $MgCl_2$ at concentrations of (A) 40 nM HMfA and (B) 7.5 nM HMfB reveal an $Mg^{2+}$ concentration-dependent increase of archaeal nucleosome formation

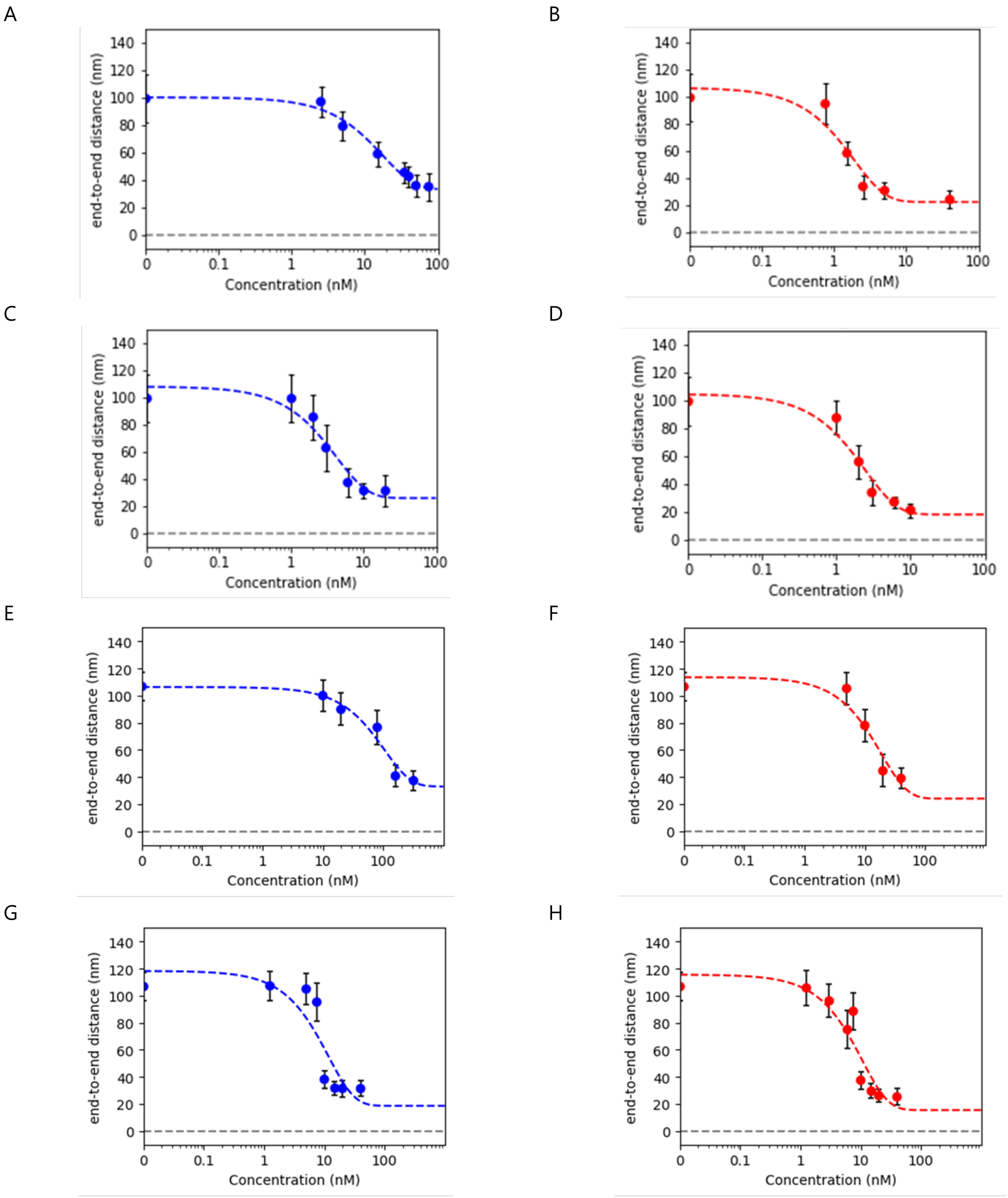


**Supplementary Figure 4. Calculated end-to-end distance as a function of protein concentration.** The EED was calculated for each concentration in the titration range. Subsequently we fitted a sigmoidal function through the data points. From the fit we are able to extract the end value and error. A, C, E, G) EED of HTkA, HTkB, HMfA and HMfB respectively. B, D, F, H) EED of HTkA, HTkB, HMfA and HMfB in the presence of 5 mM of $MgCl_2$ respectively.

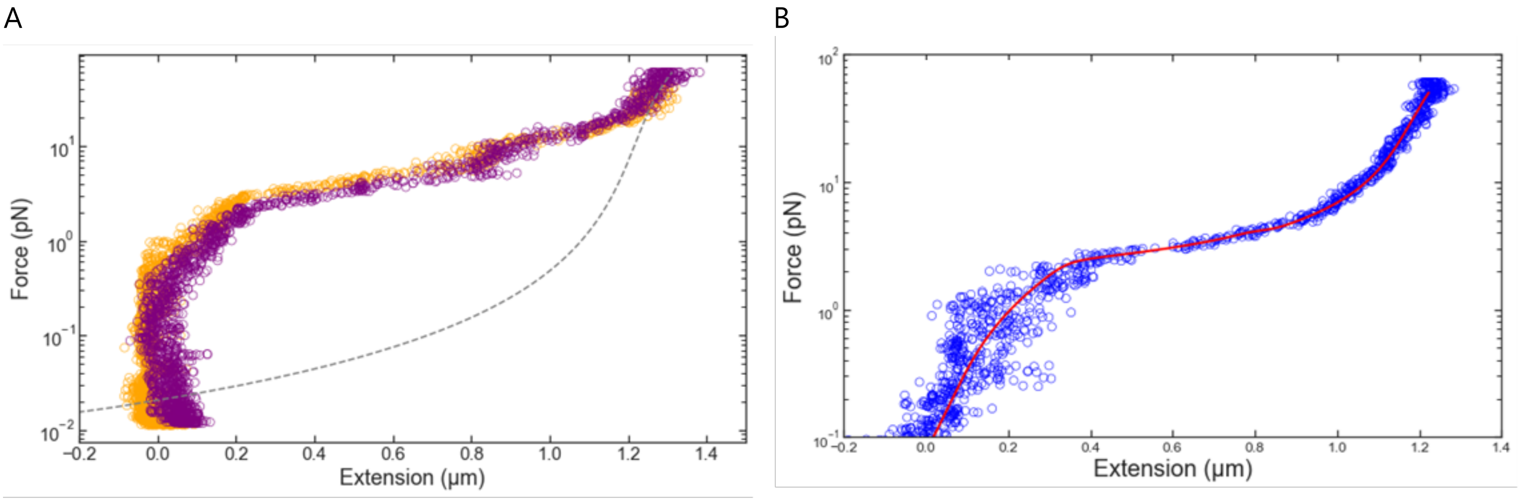


**Supplementary Figure 5.** A) A typical force-extension curve, where a HTkB hypernucleosome is being pulled (orange) and next the force is slowly released, which is reflected by the retraction curve (purple). No hysteresis is observed, indicating that the hypernucleosome complex is in thermodynamic equilibrium and that the histone dimers do not dissociate from the DNA. Hypernucleosomes that are not in thermodynamic equilibrium exhibit hysteresis and cannot be quantitatively analyzed using the model devised by Henneman et al. (2021) B) A typical force-extension curve of a HMfB hypernucleosome. The red line shows the model fitted to our data in blue. This allows us to extract quantitative values from our data.

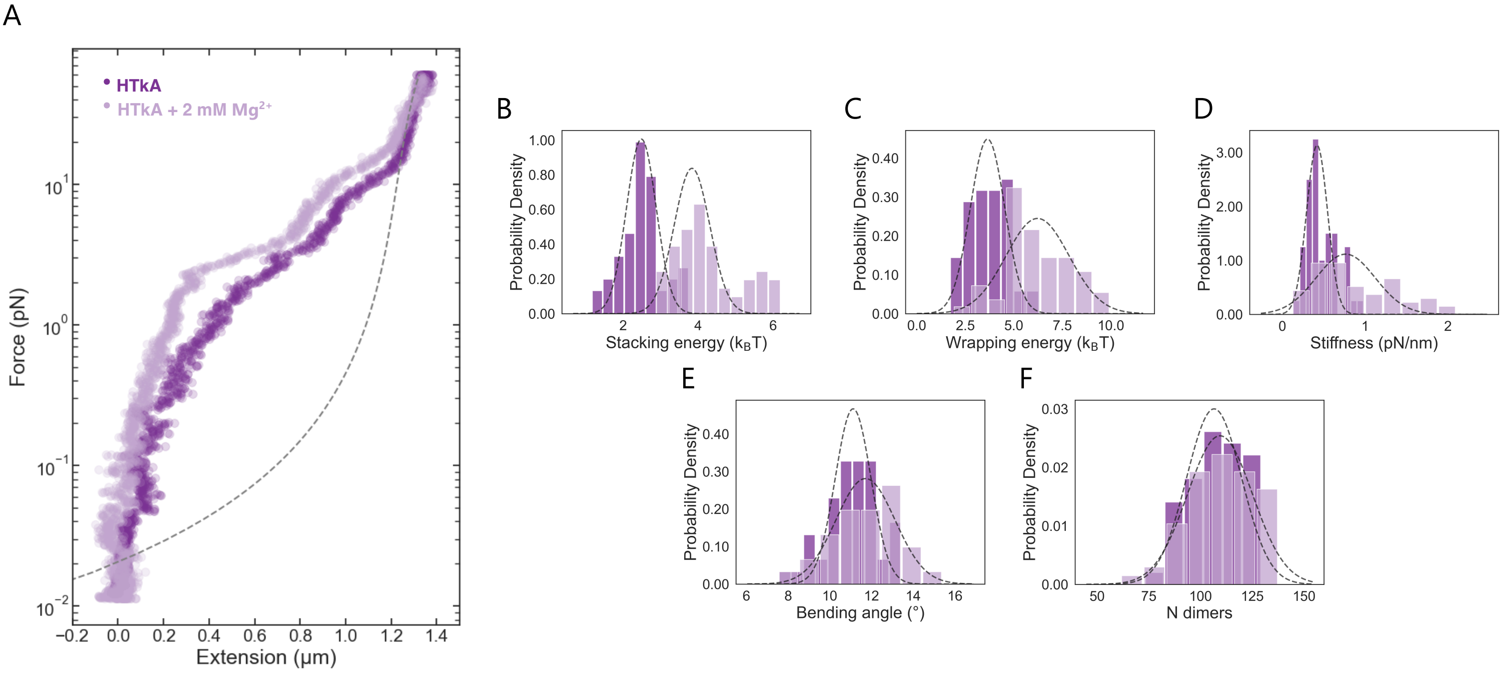


**Supplementary Figure 6.** A) Force extension curves of HTkA in the absence and the presence of 2 mM $Mg^{2+}$, as indicated. B-F) The histograms of stacking energy, wrapping energy, stiffness, bending angle, and number of bound dimers of HTkA in the absence and the presence of $Mg^{2+}$ The histograms were fitted with a normal distribution. The mean and standard error of the mean of all fit parameters can be found in Table 2.

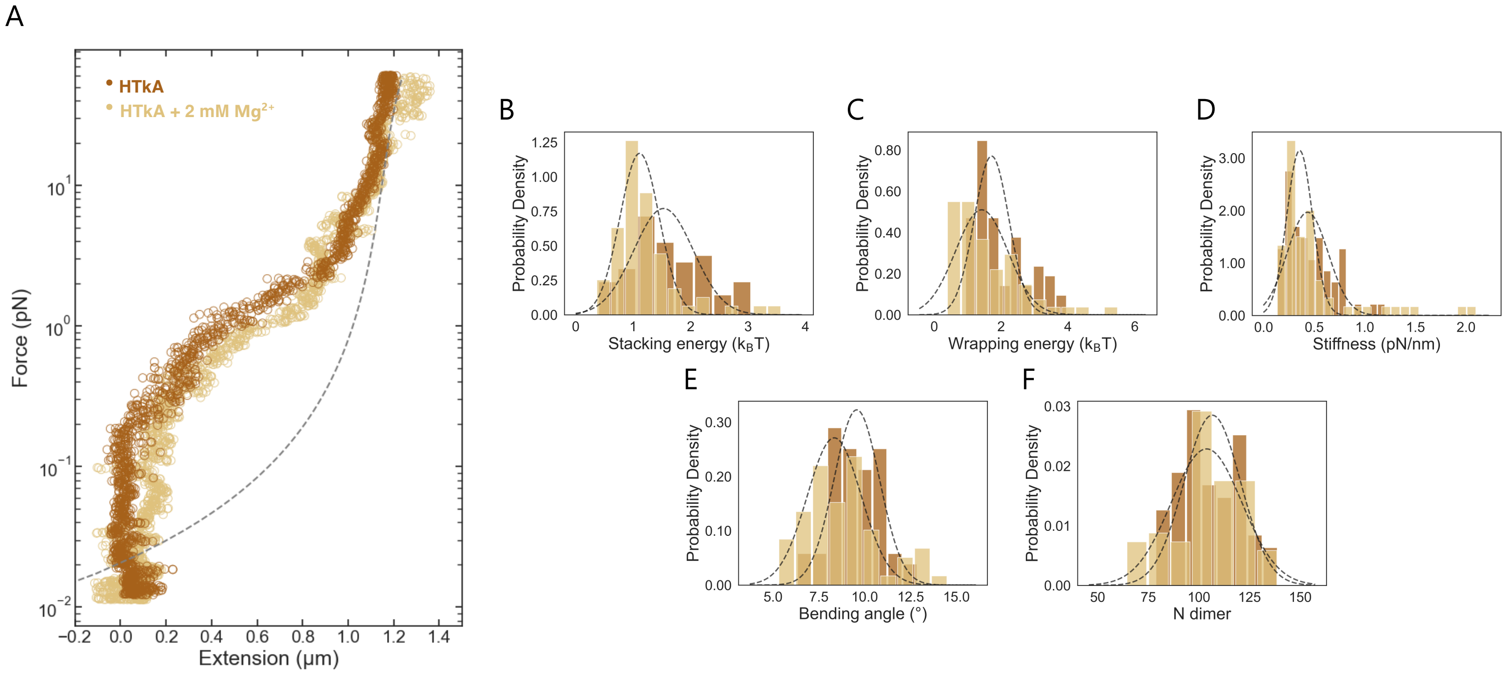


**Supplementary Figure 7.** A) Force extension curves of HMfA in the absence and the presence of 2 mM $Mg^{2+}$, as indicated. B-F) The histograms of stacking energy, wrapping energy, stiffness, bending angle, and number of bound dimers of HMfA in the absence and the presence of $Mg^{2+}$ The histograms were fitted with a normal distribution. The mean and standard error of the mean of all fit parameters can be found in Table 2.
